# A learning experience elicits sex-dependent neurogenomic responses in *Bicyclus anynana* butterflies

**DOI:** 10.1101/2022.07.27.501636

**Authors:** David A. Ernst, Gabrielle A. Agcaoili, Abbigail N. Merrill, Erica L. Westerman

## Abstract

Sexually dimorphic behavior is pervasive across animals, with males and females exhibiting different mate selection, parental care, foraging, dispersal, and territorial strategies. However, the genetic underpinnings of sexually dimorphic behaviors are poorly understood. Here we investigate gene networks and expression patterns associated with sexually dimorphic imprinting-like learning in the butterfly *Bicyclus anynana*. In this species, both males and females learn visual preferences, but learn preferences for different traits and use different signals as salient, unconditioned cues. To identify genes and gene networks associated with this behavior, we examined gene expression profiles of the brains and eyes of male and female butterflies immediately post training and compared them to the same tissues of naïve individuals. We found more differentially expressed genes and a greater number of significant gene networks in the eye, indicating a role of the peripheral nervous system in visual imprinting-like learning. Females had higher chemoreceptor expression levels than males, supporting the hypothesized sexual dimorphic use of chemical cues during the learning process. In addition, genes that influence *B. anynana* wing patterns (sexual ornaments), such as *invected*, *spalt*, and *apterous*, were also differentially expressed in the brain and eye, suggesting that these genes may influence both sexual ornaments and the preferences for these ornaments. Our results indicate dynamic and sex-specific responses to social scenario in both the peripheral and central nervous systems and highlight the potential role of wing patterning genes in mate preference and learning across the Lepidoptera.

## Introduction

Sexually dimorphic behavior is pervasive across animal taxa. Males and females may exhibit different mate selection strategies (Byrne and Rice, 2006; Kokko and Johnstone, 2002; Talyn and Dowse, 2004), parental care behavior (Trivers, 1972; Zilkha et al., 2017), foraging strategies (Ehl et al., 2018; Quillfeldt et al., 2011; Shannon et al., 2006), dispersal (reviewed in (Greenwood, 1980; Trochet et al., 2016)), and territorial displays (Reedy et al., 2017; Rosell and Thomsen, 2006). Though pervasive across species and context, the genetic underpinnings of many types of sexually dimorphic behavior are poorly understood. This is partially because males and females carry much of the same genetic material; thus, sex-specific behavior is unlikely to be allele dependent, except for the rare behaviors that are primarily associated with genes of large effect on the sex chromosome. And, because behaviors are notoriously complex traits, even sexually dimorphic behaviors influenced by genes of large effect on the sex chromosome are likely to also be influenced by autosomal genes of minor effect (Edwards et al., 2009; Lande, 1980).

Substantial headway has been made in elucidating the hormones and genes that act as master regulators of sexually dimorphic traits and behaviors in model systems. Sex-specific steroid hormone production is associated with sexually dimorphic behaviors such as song production in song birds (Alward et al., 2013; Gurney and Konishi, 1980), aggression in mammals (reviewed in (Hashikawa et al., 2018)), and spawning in fish (Pradhan and Olsson, 2015). Similarly, sex-specific alternative splicing of master regulator genes, such as *doublesex*, is associated with sexually dimorphic morphology and behavior in arthropods (Kunte et al., 2014; Rideout et al., 2007; Rodriguez-Caro et al., 2021; Wang et al., 2020). However, hormones and genes such as *doublesex* are often upstream master regulators, and the presumably sexually dimorphic downstream gene networks associated with hormone- and *doublesex*-related behaviors remain largely unknown, outside of courtship initiation in the fruit fly *Drosophila melanogaster* (Datta et al., 2008; Ruta et al., 2010) and song production in the zebra finch *Taeniopygia guttata* (Olson et al., 2015; Woodgate et al., 2014) and the canary *Serinus canaria* (Alward et al., 2018).

One sexually dimorphic behavior that is pervasive across animals is imprinting- like mate preference learning. In imprinting-like mate preference learning, sexually immature, or juvenile, individuals learn preferences for characteristics of adults (often, but not always parents) of the opposite sex (Immelmann, 1975; ten Cate and Vos, 1999; Verzijden et al., 2012). This behavior is inherently sexually dimorphic, as females learn preferences for male traits, and males learn preferences for female traits (Kendrick et al., 2001; ten Cate, 1985; Verzijden et al., 2008; Witte and Sawka, 2003). The sexual dimorphism in trait learning can be quite extreme if adults are highly sexually dimorphic or there are sex-specific signal modalities, such as male-limited pheromones or song.

To better understand the gene networks underlying sexual dimorphism in imprinting-like learning, we examined sex-specific gene expression patterns in the brains and eyes of *Bicyclus anynana* butterflies during an imprinting-like learning event. Both male and female *B. anynana* butterflies exhibit imprinting-like learning, but they learn preferences for different traits. Female *B. anynana* learn preferences for numbers of dorsal forewing eyespots and are better at learning preferences for increasing numbers of spots (Westerman et al., 2012). Conversely, male *B. anynana* learn preferences for dorsal hindwing eyespots and are better at learning preferences for loss of spots (Westerman et al., 2014). In addition to the observed sexual dimorphism in traits learned and directionality of learning bias, females learn from males who exude a volatile sex pheromone (Nieberding et al., 2008; Nieberding et al., 2012; Westerman and Monteiro, 2013), while males learn from females who, to our knowledge, do not have a volatile sex pheromone. Thus, the two sexes are likely using different cues as unconditioned stimuli to induce imprinting-like learning.

This sexual dimorphism in learning could be associated with sexual dimorphism in perception, sexual dimorphism in downstream neural processing, or a combination of these two processes. Previous studies suggest that male *B. anynana* have larger eyes and more facets (ommatidia) than female *B. anynana*, and consequently, they potentially have greater spatial acuity (Everett et al., 2012; Macias-Muñoz et al., 2015). If the observed sexual dimorphism in learning is primarily associated with sexual dimorphism in visual perception, we expect to see differential gene expression in the eyes of female and male butterflies and in visual processing genes in the brain. Alternatively, the observed sexual dimorphism in learning could be associated with sex-specific downstream processing, as is seen in *D. melanogaster*’s response to pheromones (Datta et al., 2008; Ruta et al., 2010). In this case we expect to find differential expression of genes unrelated to visual processing in the brains of males and females. We might also find differential expression of putative “magic genes,” genes subject to divergent selection that also pleiotropically affect reproductive isolation, potentially by being associated with both the production of and preference for given a trait (Servedio et al., 2011), such as butterfly wing patterning genes. Many wing patterning genes are expressed in the heads of *B. anynana* (Ernst and Westerman, 2021), and males and females have different wing patterns, with males having brighter UV-reflective eyespots than females (Everett et al., 2012; Prudic et al., 2011) while females have more dorsal hindwing spots than males (Westerman et al., 2014). Additionally, since males but not females produce pheromones that can act as the unconditioned stimuli for learning (Nieberding et al., 2008; Westerman and Monteiro, 2013), we may identify female-specific expression of genes in chemosensory processing pathways.

## Results

To examine sex-specific gene expression in the brains and eyes of *Bicyclus anynana* butterflies during an imprinting-like learning event, both male and female *B. anynana* butterflies were either subjected to an imprinting-like learning event with a conspecific of the opposite sex bearing modified wing ornaments or were placed in a cage alone as a control (Fig. 1A). These two treatments mirror the experiences of trained and naïve individuals prior to mate choice assays in published butterfly imprinting-like learning studies (Westerman et al., 2012; Westerman and Monteiro, 2013; Westerman et al., 2014). Butterflies were observed during these training/control periods, and the corresponding behavioral data were analyzed to confirm that sex-specific expression patterns were not the result of sexually dimorphic activity levels (S1 Table & S2 Table). We then sequenced the eye and brain transcriptomes of these animals, N=10 per treatment per sex, which generated a total of nearly three billion high-quality 50 base pair (bp) single end (SE) reads (S3 Table). Approximately 1.6 million reads (0.05% of raw reads) were removed during adapter trimming, with 2.7 billion of the remaining reads (90% of trimmed reads) mapping to the *B. anynana* reference genome (Nowell et al., 2017). Across all brain libraries, 16,785 genes (74% of annotated genes in the genome) had at least 10 mapped reads, while this was the case for 16,612 genes (73%) for eye libraries. For each tissue, these gene sets were used as input for differential expression analyses.

**Figure 1:**
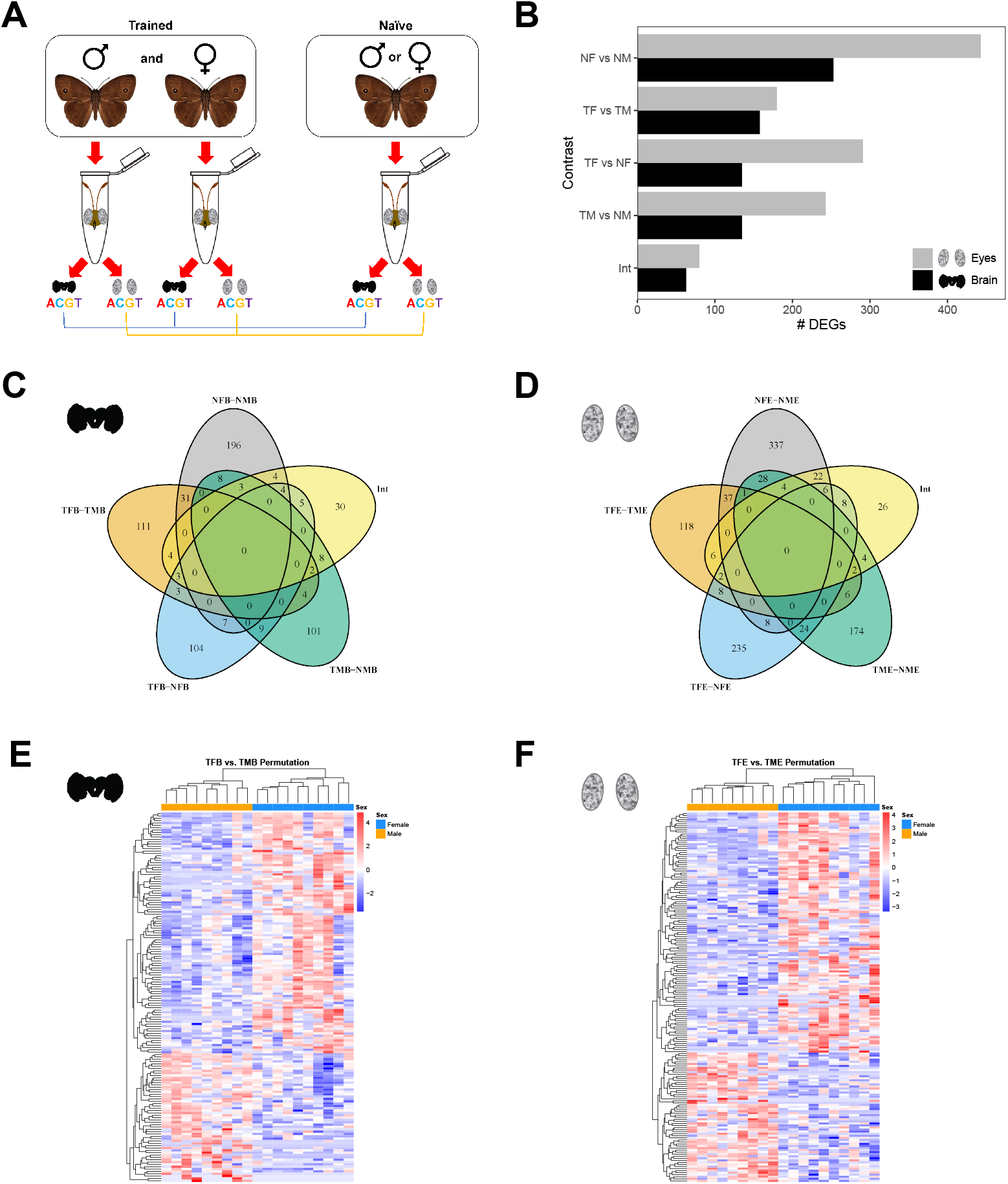
Experimental design and broadscale sexually dimorphic gene expression. A) Protocol for butterfly training and sampling. Newly emerged males/females were either solo or paired with a two-day-old, zero-spot female/four-spot male. Heads of each focal animal were collected, the brain and eyes dissected, and mRNA sequenced for expression analysis. B) Numbers of differentially expressed genes for each comparison for each tissue. C) Brain Venn diagrams showing overlap patterns for differentially expressed genes. D) Eye Venn diagrams showing overlap patterns for differentially expressed genes. E) Brain gene expression heatmaps of differentially expressed genes from trained females vs. trained males. Each row indicates a single gene, and each column indicates an individual sample. Counts were first normalized by variance stabilizing transformation, and gene-wise Z-scores were calculated for plotting. Genes and samples are clustered by expression, with warmer colors denoting increased expression relative to the mean for a given gene, while cooler colors denote decreased expression relative to the mean. F) Eye gene expression heatmaps of differentially expressed genes from trained females vs. trained males. NFB=naïve female brain, NMB=naïve male brain, TFB=trained female brain, TMB=trained male brain, NFE=naïve female eye, NME=naïve male eye, TFE=trained female eye, TME=trained male eye, Int=interaction.

During data quality assessment, gene expression clustering analysis revealed that one sample (TMB_E2, a trained male brain sample) was likely mislabeled, as it clustered with eye samples (S1 Fig.). Because the two tissue types exhibited distinct clustering patterns and tissue type accounted for approximately 85% of the variance, this sample was discarded and not included in downstream analyses.

For all differential gene expression comparisons, we used DESeq2 to perform both a standard differential expression analysis as well as a permutation-test-based analysis, a method that eliminates the assumption of gene independence and provides a more accurate representation of the data structure of gene expression datasets (Bloch et al., 2018; Ghalambor et al., 2015; Slonim, 2002). Nearly all genes that were determined to be differentially expressed in the standard DESeq2 analyses (Tables S4-S15) were also identified as differentially expressed when employing permutation test analyses (Tables S4-S15). Moreover, because the permutation test analyses reduce potential over-correction by multiple testing correction methods, a larger number of differentially expressed genes (DEGs) was found for all comparisons. Therefore, all downstream analyses were conducted with the results of the permutation-based differential expression tests. While all DEG sets obtained from these analyses were tested for gene ontology (GO) term enrichment, GO term enrichment results are only reported for DEG sets with significantly enriched GO terms.

### Trained male and female brains have distinct expression patterns

Contrasting naïve female and male brains revealed a baseline of 253 genes that were differentially expressed (Fig. 1B,C; S4 Table). Conversely, 158 genes were found to be differentially expressed between trained female and male brains (Fig. 1B,C,E; S5 Table). Of these gene sets, 127 genes were unique to the training contrast (Fig. 1C), several of which are linked to various neural processes, including neurodevelopment, neural signaling, eye development, and phototransduction (Fig. 2; S16 Table). Additionally, four genes with putative chemosensory functions were differentially expressed, all of which were upregulated in females relative to males (chemosensory protein 6, *BANY.1.2.g12995*; ejaculatory bulb-specific protein 3-like, *BANY.1.2.g12992*; ejaculatory bulb-specific protein 3-like, *BANY.1.2.g12993*; and odorant receptor Or2-like, *BANY.1.2.g25738*) (Ernst and Westerman, 2021). Finally, a gene encoding vitellogenin-like (*BANY.1.2.g11921*), a protein known to influence the social behavior of numerous insect species (Morandin et al., 2019; Nelson et al., 2007; Roy-Zokan et al., 2015), was also upregulated in females.

**Figure 2:**
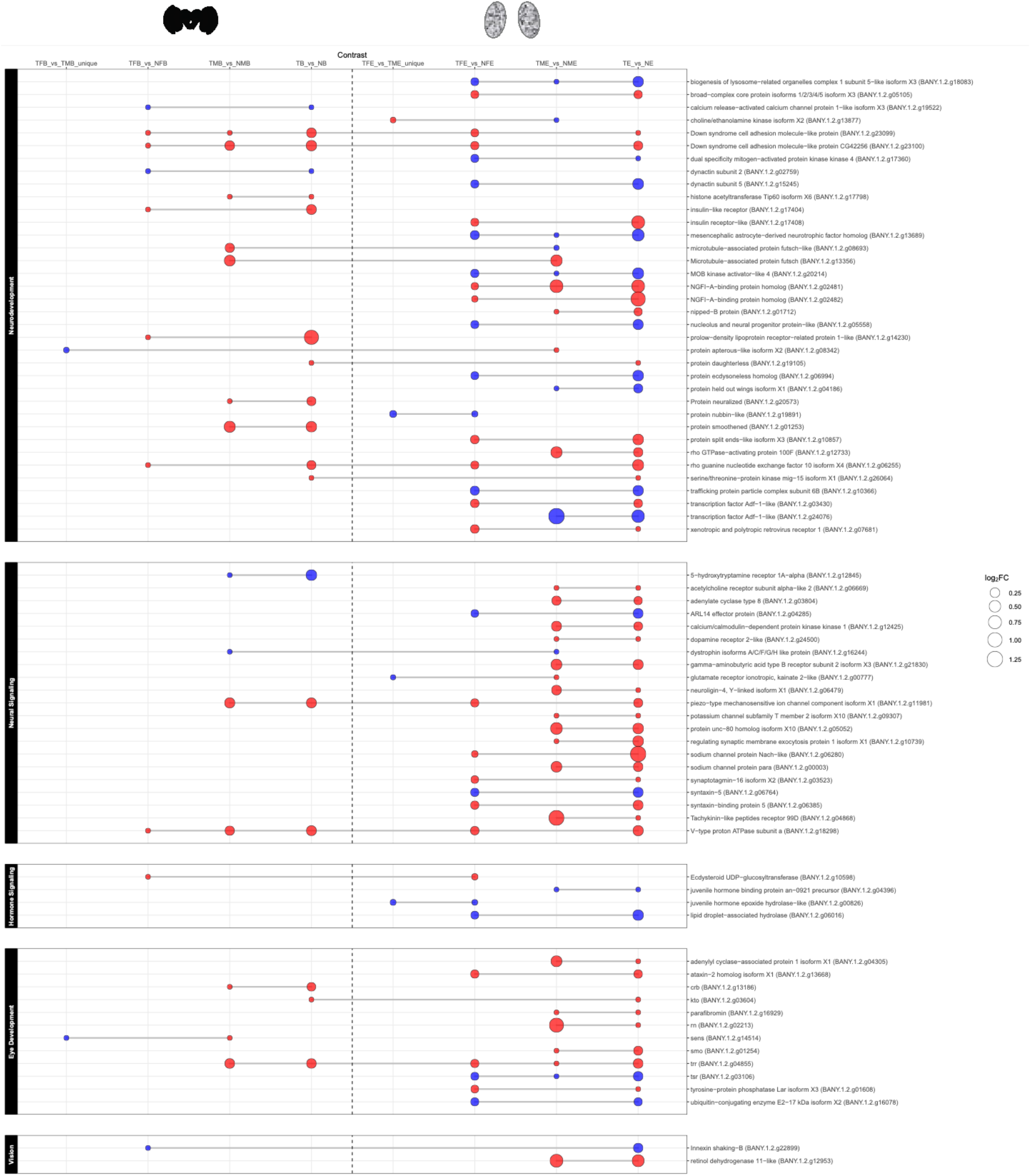
Neural processing, hormone signaling, and vision genes are differentially expressed in multiple contrasts. The size of each dot indicates the effect size (log_2_FC), while the color indicates the gene regulation relative to the first sample type listed for the contrast (e.g., for the TB vs. NB contrast, red indicates upregulation in trained brains, and blue indicates downregulation in trained brains). Gray lines connecting the dots denote that the gene was differentially expressed across multiple contrasts. NFB=naïve female brain, NMB=naïve male brain, TFB=trained female brain, TMB=trained male brain, NFE=naïve female eye, NME=naïve male eye, TFE=trained female eye, TME=trained male eye, TB=trained brain, TE=trained eye, NB=naïve brain, NE=naïve eye.

GOExpress analyses, which find gene ontology (GO) terms that best classify samples from two separate groups, identified 171 GO terms that were significantly associated with differences between naïve female and male brains (p < 0.05; S17 Table), while 166 GO terms differentiated trained female and male brains (p < 0.05; S18 Table). To eliminate baseline differences, we removed significant terms that were also found in the naïve results, resulting in 51 GO terms linked to differences specific to training (S18 Table). Of these terms, several are linked to neural processing, including calmodulin binding (p = 0.004), vesicle docking involved in exocytosis (p = 0.042), gap junction (p = 0.046), and neuropeptide signaling pathway (p = 0.008).

### Trained male and female eyes have distinct expression patterns

Differential expression analysis for naïve female and male eyes found a baseline of 443 genes that were differentially expressed between naïve female and male eyes (Fig. 1B,D; S6 Table). By contrast, 180 DEGs were found for the trained female vs. male comparison (Fig. 1B,D,F; S7 Table). In total, 142 genes were unique to the trained eye contrast (Fig. 1D), including genes encoding proteins linked to neurodevelopment, neural signaling, hormone signaling, and vision (Fig. 2; S16 Table). Moreover, three genes putatively linked to circadian rhythms showed differential expression, including circadian clock-controlled protein-like (*BANY.1.2.g04378*), which was upregulated in males, and circadian clock-controlled protein-like (*BANY.1.2.g05915*) and protein takeout-like (*BANY.1.2.g05914*), which were both upregulated in females. The takeout gene (*to*) is also associated with male courtship behavior in *D. melanogaster* (Dauwalder et al., 2002).

GOExpress analyses revealed 165 and 138 GO terms that were significantly linked to expression differences between the sexes for naïve and trained eyes, respectively (p < 0.05; S19, S20 Tables). Removal of terms that overlapped both the naïve and trained sets resulted in 37 GO terms linked to sex-specific differences in response to training (S20 Table). A number of these terms were associated with neural processes and sensory transduction, including chloride transmembrane transport (p = 0.007), chloride channel activity (p = 0.01), vesicle docking involved in exocytosis (p = 0.017), and G protein-coupled peptide receptor activity (p = 0.025).

### Training has sex-dependent effects on expression patterns in brains and eyes

Sex-specific pairwise comparisons between trained and naïve tissues revealed many DEGs in all sex-dependent comparisons.

Starting with the female comparisons, a total of 135 genes were found to be differentially expressed between trained and naïve female brains (Fig. 1B,C; S8 Table), many of which have potential roles in neural development, neural signaling, hormone metabolism, and eye-related processes (Fig. 2; S16 Table).

For the trained vs. naïve female eyes comparison, differential expression analysis identified 291 DEGs (Fig. 1B,D; S9 Table). GO enrichment analysis found 12 GO terms enriched in this gene set, with the top being mitochondrion (FDR=4.04E-04), intracellular organelle (FDR=4.04E-04), and organelle (FDR=5.23E-04) (S21 Table). There were several genes of interest in the trained vs. naïve female eye contrast, including genes linked to neural development and signaling, hormone signaling, eye development, and vision (Fig. 2; S16 Table).

Similar to the female brains comparison, the trained vs. naïve male brains comparison found 135 DEGs (Fig. 1B,C; S10 Table), including several genes associated with neurodevelopment, neural signaling, and eye development (Fig. 2; S16 Table).

Differential expression analysis revealed 243 DEGs for the trained vs. naïve male eyes comparison (Fig. 1B,D; S11 Table). Again, numerous genes involved with neural development, neural signaling, hormone signaling, vision, and eye development were found to be differentially expressed between trained and naïve male eyes (Fig. 2; S16 Table).

Moreover, 63 genes in the brain and 80 genes in the eye were found to have a significant sex:condition interaction, indicating that training differentially affected their expression in females versus males (Fig. 1B,C,D; S12, S13 Tables). In both tissues, these sex:condition interactions were found for genes involved with neural development and signaling, and interactions were also found for genes linked to eye development in the eye comparison (S16 Table). In addition, a gene putatively involved with chemoreception (olfactory receptor 21, *BANY.1.2.g12009*; (Ernst and Westerman, 2021)) and a gene associated with regulating circadian rhythms (protein LSM12 homolog, *BANY.1.2.g13734*; (Lee et al., 2017)) showed significant sex:condition interactions in the brain and eyes, respectively.

### Training has a sex-independent effect on gene expression in brains

Testing for the overall effect of training while controlling for differences in expression due to sex revealed 283 genes that were differentially expressed in trained vs. naïve brains (Fig. S2A; S14 Table). Many of the genes in this gene set have functions related to neurodevelopment, neural signaling, hormone signaling, and eye development (Fig. 2; S16 Table). Moreover, LSM12 homolog (*BANY.1.2.g13734*), which showed significant sex:condition interactions in the eyes, was also differentially expressed and was upregulated in naïve brains.

### Training has a sex-independent effect on gene expression in eyes

In total, 658 DEGs were identified for the trained vs. naïve eyes comparison when controlling for sex (Fig. S2B; S15 Table). GO enrichment analysis revealed 30 enriched GO terms, with the top terms being mitochondrion (FDR=1.92E-06), protein-containing complex (FDR=3.03E-06), and intracellular organelle (FDR=5.17E-06) (S22 Table).

Several of these DEGs have putative functions in neurodevelopment, neural signaling, hormone signaling, eye development, and vision (Fig. 2; S16 Table). In addition, a number of genes linked to learning and memory were differentially expressed between trained and naïve eyes. Several of these genes were upregulated in trained eyes, including nipped-B protein (*BANY.1.2.g01712*), Ca(2+)/calmodulin-responsive adenylate cyclase (*BANY.1.2.g01825*), two transcription factor Adf-1-like (*BANY.1.2.g03430* and *BANY.1.2.g08959*), adenylate cyclase type 8 (*BANY.1.2.g03804*), neurobeachin-like (*BANY.1.2.g12252* and *BANY.1.2.g12258*), and ataxin-2 homolog isoform X1 (*BANY.1.2.g13668*). Conversely, cyclic AMP response element-binding protein B isoform X3 (*BANY.1.2.g01685*), one transcription factor Adf-1-like (*BANY.1.2.g24076),* probable RNA helicase armi isoform X1 (*BANY.1.2.g17424*), and fatty acid-binding protein-like (*BANY.1.2.g17524*) were upregulated in naïve eyes. Finally, two genes involved with male courtship in *Drosophila* (calcium/calmodulin-dependent 3’,5’-cyclic nucleotide phosphodiesterase 1 isoform X1, *BANY.1.2.g07806*; and cytoplasmic dynein 2 heavy chain 1, *BANY.1.2.g19627*) were upregulated in *B. anynana* eyes in the training condition.

### One gene network is associated with training condition in the brain

To investigate gene networks that are associated with an imprinting-like learning experience, we performed tissue-specific weighted gene co-expression network analyses (WGCNA). Brain co-expression network analysis identified 17 modules, which was reduced to 11 modules after merging highly correlated modules (Fig. 3A; S3A Fig.). Of these modules, only one (the red module) was significantly correlated with a trait, specifically the trained male brain vs. naïve male brain contrast (i.e., the red module was significantly correlated with training condition for male brains; r=0.6; FDR=0.004) (Fig. 3C; S3B Fig.). This module consisted of 655 genes (S23 Table), with the top hub gene (i.e., the most highly connected gene) identified as NADH dehydrogenase [ubiquinone] 1 beta subcomplex subunit 7-like (*BANY.1.2.g00209*). GO enrichment analyses found five significantly enriched GO terms in the red module, which were linked to nucleic acid and cyclic compound binding and mRNA metabolism (S24 Table; Fig. 4A).

**Figure 3:**
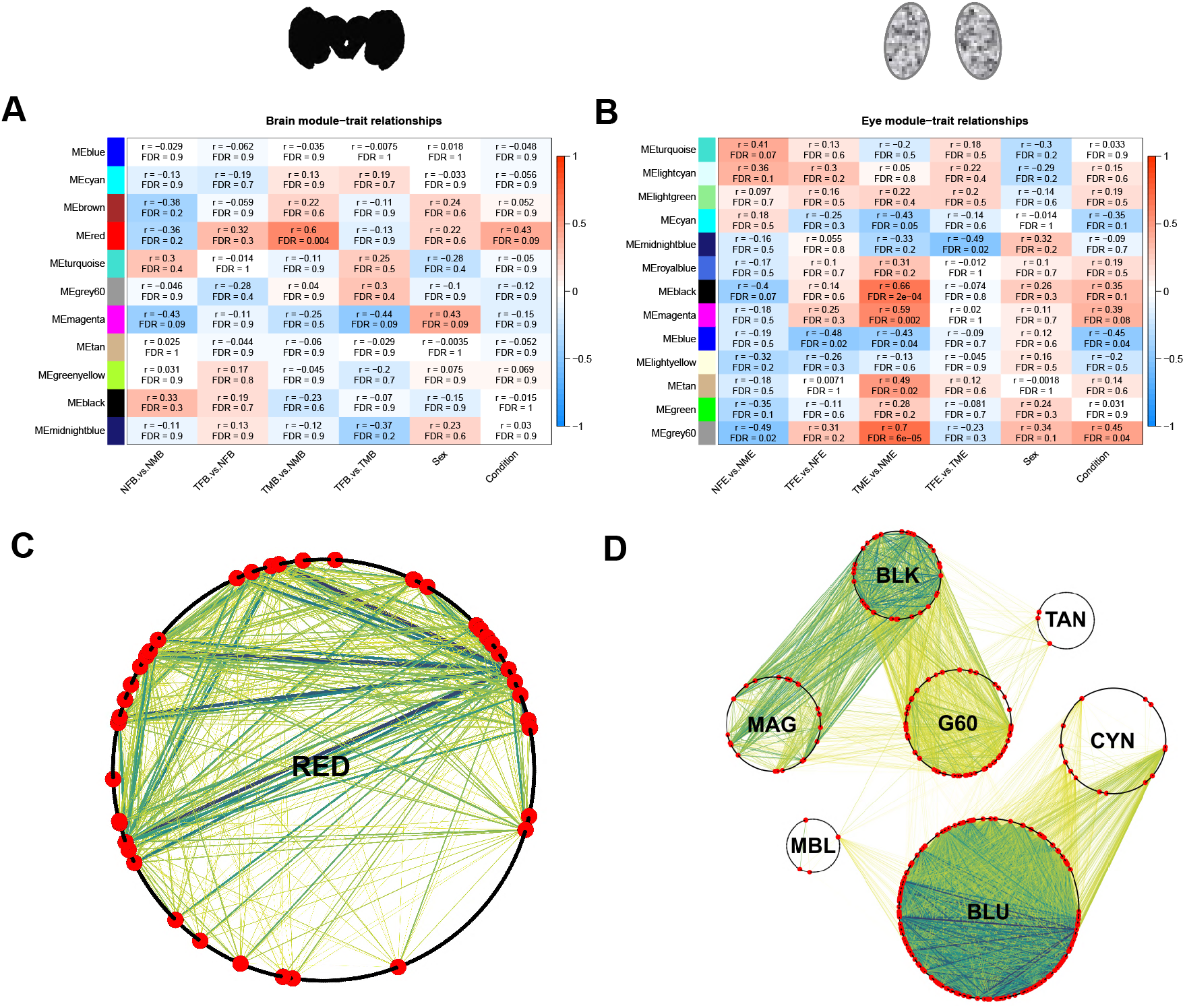
Gene network modules in brain and eyes are significantly associated with training. Significant modules from co-expression analyses. A) Brain module-trait association heatmap. Rows indicate module eigengenes (ME), and columns indicate the pairwise binary indicators representing the various comparisons (“traits”) of interest. The top numbers in each cell denote the correlation value (r), with false discovery rate (FDR) values below. Cells are colored by the strength of the association, with r ranging from -1 to 1. B) Eye module-trait association heatmap. C) WGCNA brain analysis red module Cytoscape plot. Each black dot around the perimeter of the circle indicates a node (gene), with larger red dots indicating differentially expressed genes from the contrast for which the module is significantly associated (i.e., trained vs. naïve male brain). Each line indicates an edge (connection) for differentially expressed genes within the module, with thinner yellow lines indicating weaker connections and thicker blue lines indicating stronger connections. D) WGCNA eye analysis, Cytoscape plot of all significant modules. Only edges for differentially expressed genes within and between modules are shown. BLK=black module, BLU=blue module, CYN=cyan module, G60=grey60 module, MAG=magenta module, MBL=midnightblue module, RED=red module, and TAN=tan module.

**Figure 4:**
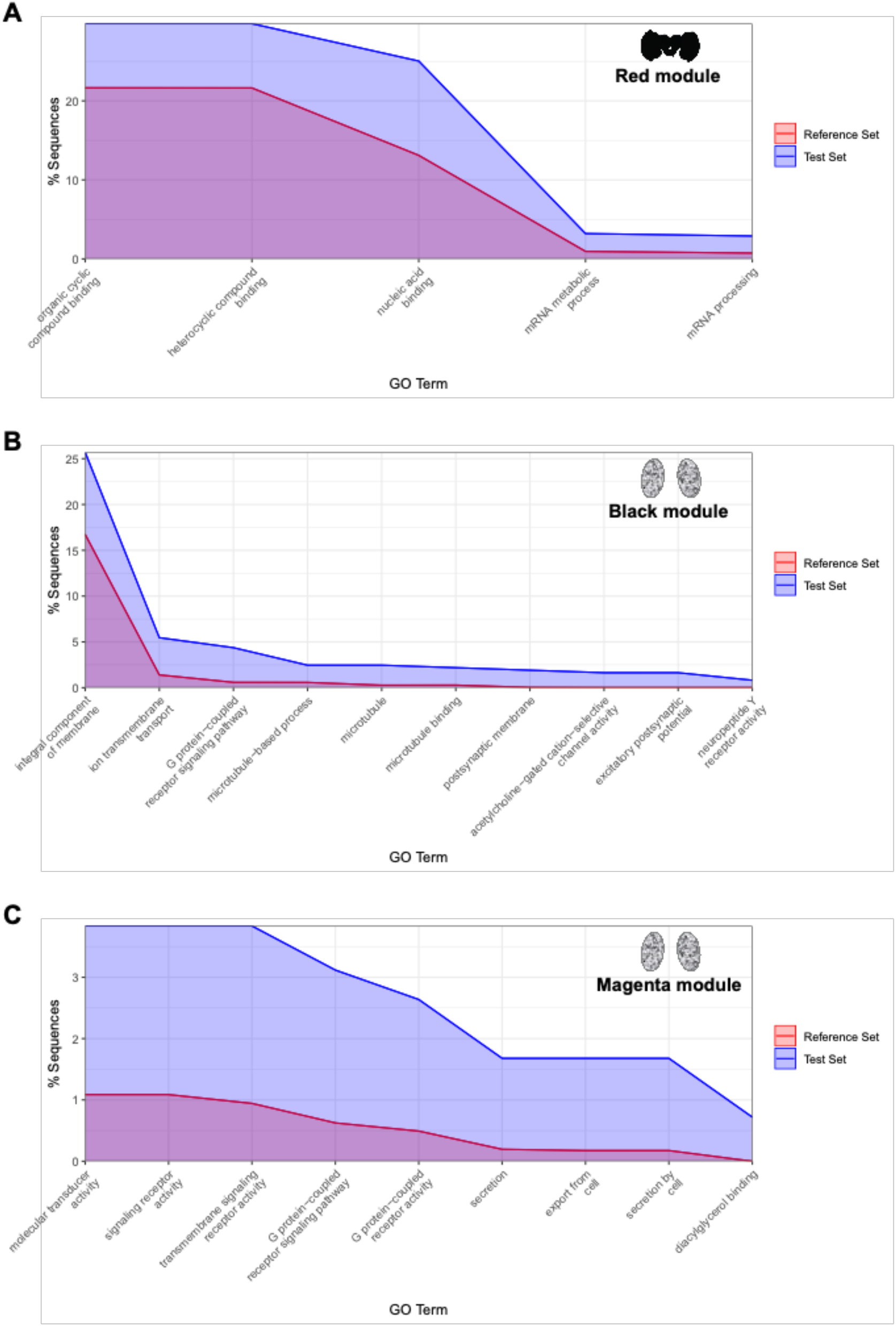
Gene ontology enrichment plots for significant brain and eye modules of interest. A) Significantly enriched GO terms in the brain red module. For each GO term, the percentage of sequences annotated with that term within the Test Set (i.e., all red module genes) is plotted along with the percentage of sequences annotated with that term within the Reference Set (i.e., all genes used in the co-expression analysis). B) Significantly enriched GO terms in the eye black module. Due to the large number of enriched GO terms in this module, only the most specific terms identified by Blast2GO were plotted for clarity. C) Significantly enriched GO terms in the eye magenta module.

Many genes within the red module are linked to various neural and sensory processes. Of particular interest, 41 DEGs identified in the trained vs. naïve male brain contrast were also present in the red module network (S23 Table). Many of these genes encode proteins linked to neural development, such as protein smoothened (*BANY.1.2.g01253*), protein abrupt-like isoform X5 (*BANY.1.2.g17381*), histone acetyltransferase Tip60 isoform X6 (*BANY.1.2.g17798*), Down syndrome cell adhesion molecule-like protein (*BANY.1.2.g23099*), Down syndrome cell adhesion molecule-like protein CG42256 (*BANY.1.2.g23100*), and helicase domino (*BANY.1.2.g24509*). Additionally, others encode proteins involved with neural signaling, such as piezo-type mechanosensitive ion channel component isoform X1 (*BANY.1.2.g11981*) and V-type proton ATPase subunit a (*BANY.1.2.g18298*) and eye development, such as *trr*, (*BANY.1.2.g04855*) and *crb* (*BANY.1.2.g13186*) (S16 Table). In addition to its role in eye development, *trr* is also involved with short term courtship memory in *D. melanogaster* (Sedkov et al., 2003).

### Several gene networks are associated with training condition in the eyes

Eye co-expression network analysis identified 20 modules, which was reduced to 13 modules after merging highly correlated modules (Fig. 3B, S4A Fig.). Of these modules, seven (the black, blue, cyan, grey60, magenta, midnight blue, and tan modules) were significantly correlated with at least one contrast, and DEGs for the correlated contrast(s) were present in all seven of these modules (Fig. 3D, S4 Fig.; Tables S25-S37). Two of these modules (the black and magenta modules), both of which were significantly correlated with the trained male vs. naïve male eyes contrast, were of particular interest based on their GO enrichment profiles. The black module (r=0.66; FDR=2E-04) consisted of 366 genes centered around the top hub gene gamma-aminobutyric acid type B receptor subunit 2 (*BANY.1.2.g00039*), a component of the receptor for the neurotransmitter GABA (S27 Table; Fig. 3D, S4C Fig.). Moreover, 73 GO terms were enriched in this module, most of which are associated with neural processes (e.g., neurotransmitter receptor activity involved in regulation of postsynaptic membrane potential, chemical synaptic transmission, and excitatory postsynaptic potential) (S28 Table; Fig. 4B). A total of 32 DEGs from the trained male vs. naïve male eyes contrast were found in the black module, nearly half of which are associated with neural and eye development and neural signaling. Differentially expressed development genes include protein unc-80 homolog isoform X10 (*BANY.1.2.g05052*), microtubule-associated protein futsch-like (*BANY.1.2.g08693*), delta and Notch-like epidermal growth factor-related receptor (*BANY.1.2.g09881*), protein abrupt-like isoform X1 (*BANY.1.2.g17383*), and *rst* (*BANY.1.2.g15359*) (S16 Table; S27 Table). Moreover, differentially expressed neural signaling genes include sodium channel protein para (*BANY.1.2.g00003*), potassium voltage-gated channel subfamily KQT member 1 isoform X2 (*BANY.1.2.g01557*), adenylate cyclase type 8 (*BANY.1.2.g03804*), neuroligin-4, Y-linked isoform X1 (*BANY.1.2.g06479*), acetylcholine receptor subunit alpha-like 2 (*BANY.1.2.g06669*), potassium channel subfamily T member 2 isoform X10 (*BANY.1.2.g09307*), calcium/calmodulin-dependent protein kinase kinase 1 (*BANY.1.2.g12425*), sodium leak channel non-selective protein (*BANY.1.2.g19402*), gamma-aminobutyric acid type B receptor subunit 2 isoform X3 (*BANY.1.2.g21830*), and dopamine receptor 2-like, (*BANY.1.2.g24500*) (S16 Table; S27 Table).

The magenta module (r=0.59; FDR=0.002) consisted of 417 genes with a hub gene of disintegrin and metalloproteinase domain-containing protein 33-like (S29 Table; S4D Fig.) and showed an enriched GO term profile similar to that of the black module (S30 Table; Fig 4C). Specifically, the terms transmembrane signaling receptor activity, G protein-coupled receptor signaling pathway, G protein-coupled receptor activity, signaling receptor activity, and molecular transducer activity were found to be enriched in both the black and magenta modules. In total, 21 DEGs from the trained male vs. naïve male eyes contrast were found in the magenta module, a third of which have putative functions in neurodevelopment (protein smoothened isoform X2, *BANY.1.2.g01254*; putative defective proboscis extension response, *BANY.1.2.g12002*; rho GTPase-activating protein 100F, *BANY.1.2.g12733*; and dynamin-like 120 kDa protein, mitochondrial, *BANY.1.2.g23042*), neural signaling (regulating synaptic membrane exocytosis protein 1 isoform X1, *BANY.1.2.g10739*; and dopamine receptor 1, *BANY.1.2.g24271*), and eye development (adenylyl cyclase-associated protein 1 isoform X1, *BANY.1.2.g04305*) (S16, S29 Tables).

### Wing patterning genes are differentially expressed in both the brain and eyes

To investigate whether putative “magic genes,” or genes that influence both a given trait as well as preference for that trait, are expressed in the brain and eyes of *B. anynana*, we also explored the expression patterns of known butterfly wing patterning genes. A total of 53 wing patterning genes were found to be expressed in the brain, while 50 were expressed in the eyes (S38 Table). Although none of these wing patterning genes exhibited sex-specific expression (meaning only expressed in one sex) in either tissue, 46 were in common across the two tissues. Seven genes showed brain-specific expression, including homologs for cortex (*BANY.1.2.g04256*), engrailed (*BANY.1.2.g14935*), CD63-antigen (*BANY.1.2.g12556*), aristaless (*BANY.1.2.g21346* and *BANY.1.2.g24453*), and BarH-1 (*BANY.1.2.g19326* and *BANY.1.2.g22154*), while four exhibited eye-specific expression, including homologs for hedgehog (*BANY.1.2.g04016*) and CD63-antigen (*BANY.1.2.g20540*, *BANY.1.2.g25497*, and *BANY.1.2.g25594*).

Several wing patterning genes were identified as differentially expressed for various contrasts, including between and within sexes, in both tissue types. For the naïve female vs. male brain contrast, sal-like protein 1 (*BANY.1.2.g09547*) and CD63 antigen-like (*BANY.1.2.g23713*) are both upregulated in females (Fig. 5, Table S4). In the trained female vs. male brain contrast protein apterous-like isoform X2 (*BANY.1.2.g08342*) is upregulated in males (Fig. 5, S5 Table). In the naïve female vs. male eye contrast CD63 antigen-like (*BANY.1.2.g25497*) is upregulated in females (Fig. 5, S6 Table). Moreover, in the eye interaction contrast CD63 antigen-like (*BANY.1.2.g25497*) is upregulated in trained females and naive males (Fig. 5, S13 Table), and in the trained vs. naïve eye controlling for sex contrast CD63 antigen-like isoform X2 (*BANY.1.2.g10818*) is upregulated in naïve eyes (Fig. 5, S15 Table).

**Figure 5:**
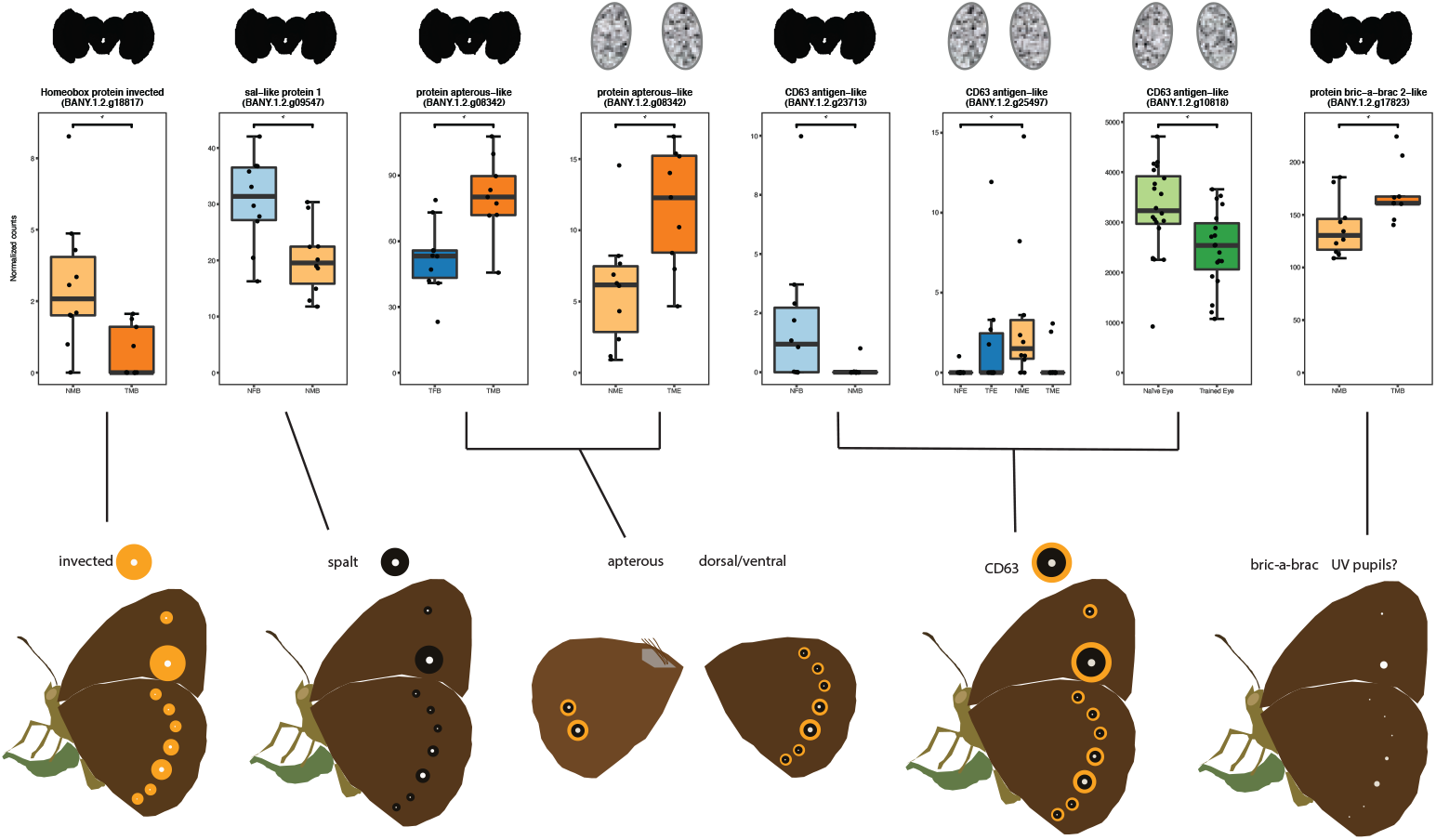
Genes that influence *B. anynana* wing patterns are also differentially expressed in the brain and eye during training. Top panel contains box plots of differentially expressed genes in different contrasts. Bottom panel indicates the elements of butterfly wing pattern (gold ring, eye spot center, black ring, whole eye spot, or dorsal/ventral identity) influenced by the corresponding differentially expressed gene. For top panel, light hue = naïve, dark hue = trained, orange = male, blue = female, green = condition (general trained/naïve).

When comparing within sexes, three known wing patterning genes were differentially expressed in male brains or eyes. In the trained vs. naïve male brain contrast, protein bric-a-brac 2-like isoform X4 (*BANY.1.2.g17823*) is upregulated in trained males while Homeobox protein invected (*BANY.1.2.g18817*) is upregulated in naïve males (Fig. 5, S10 Table). By contrast, in the trained vs. naïve male eye comparison protein apterous-like isoform X2 (*BANY.1.2.g08342*) is upregulated in trained males (Fig. 5, S11 Table). No known wing pattern genes were differentially expressed in female-specific contrasts.

## Discussion

Here we identified a number of genes that were differentially expressed in the brains and eyes of females and males during an imprinting-like learning event, as well as several associated gene networks. We found DEGs in both tissue types, suggesting that imprinting-like learning, and sexually dimorphic aspects of this learning process, are associated with transcriptional change in both the peripheral sensory system and the brain. A number of chemosensory genes were upregulated in females relative to males, supporting the hypothesized female-specific use of pheromones in the mate preference learning process (Westerman and Monteiro, 2013; Westerman et al., 2014). Furthermore, a suite of butterfly wing patterning genes, which have long been hypothesized to also influence mate preference and potentially serve as “magic genes,” were also differentially expressed in the eyes and brains of *B. anynana* butterflies during training events, further supporting their hypothesized role in mate preference and speciation.

One of the more interesting aspects of sexually dimorphic imprinting-like learning in *B. anynana* is the presence/absence of sex pheromones in males versus females. Previous studies have shown that the male sex pheromone is an indicator of age (Nieberding et al., 2012), is species-specific (Bacquet et al., 2015; Nieberding et al., 2008), is equally weighted with visual signals during female mate selection (Costanzo and Monteiro, 2007), and influences the valence females learn to associate with visual signals during imprinting-like learning (Westerman and Monteiro, 2013). Thus, male chemical cues are known to be important for female mate choice in this system. On the other hand, a sex pheromone has not been discovered in female *B. anynana*, and it remains unclear what unconditioned stimulus males use to assign positive valence to number of hindwing spots. The results of this study appear to support this sex-specific use of olfactory signals during the learning process. The most clear-cut finding supporting this hypothesis is that chemosensory genes are upregulated in females relative to males during the training period. A second result that may be related to the differential use of olfactory cues during the learning (and mate choice) process is that we found a larger set of gene networks associated with the training condition in the brains and eyes of males than in females. This could be a result of imprinting-like learning being more consistent in males than females (Westerman et al., 2012; Westerman et al., 2014).

However, it could also be a side effect of females relying more heavily on olfactory signals than males, as we did not include antennae in our analyses and consequently may have missed learning-associated gene networks that reside in female antennae. Female *Heliconius melpomene* and *Heliconius cydno* butterflies are sensitive to male pheromones (Byers et al., 2020) and exhibit different antennae expression profiles before and after copulation as well as sex-specific expression profiles (van Schooten et al., 2020). It would be interesting to see if *B. anynana* females exhibit training-specific, sexually dimorphic antennae expression profiles that correspond to their sex-specific emphasis on olfactory signals during the preference learning and mate selection process.

While the gene expression patterns of the antennae are unknown for these animals, we did find training-specific, sexually dimorphic gene expression patterns in *B. anynana* eyes. Because female and male *B. anynana* butterflies learn preferences for different visual signals and exhibit visual learning biases in different directions (gains and losses, respectively (Westerman et al., 2012; Westerman et al., 2014)), one of our hypotheses was that we would see sexually dimorphic expression of vision-related genes during the learning process, especially in the eyes. Although we did not observe differential expression of any opsins, we did find sex-dependent expression patterns of a number of vision-related genes, including an ommochrome-binding protein, retinol dehydrogenase 11, rhodopsin kinase 1 (*Gprk1*), and arrestin homolog isoform X2.

Ommochrome pigments act as filtering pigments in the eyes of butterflies, limiting the wavelengths of light a butterfly can see (Arikawa and Stavenga, 2014; Stavenga, 2002). These filtering pigments are sexually dimorphic in a number of different species, including *H. cydno*, *H. melpomene*, *Heliconius pachinus*, and *Colias erate*, and are hypothesized to influence mate choice in these systems (Buerkle et al., 2022; Ogawa et al., 2013). It remains unclear whether filtering pigment type or distribution is sexually dimorphic in *B. anynana*, or whether filtering pigment production or distribution in the eye is plastic in response to circadian rhythms, social scenario, or age. However, our findings of socially-dependent expression patterns of ommochrome-binding protein and a number of other vision-related genes suggest that vision is highly dynamic, not just in the context of light environment (Obara et al., 2008; Sakai et al., 2018; Wright et al., 2020) and circadian rhythms (Li et al., 2008; Li et al., 2005), but also in response to social environment.

In addition to finding vision-associated differentially expressed genes, a number of learning and memory genes were differentially expressed specifically in the eyes, including dopamine receptors. Moreover, the most highly connected gene for a gene network associated with training condition in male eyes (the black module) encodes a component of the receptor for the neurotransmitter GABA, gamma-aminobutyric acid type B receptor subunit 2. This network also contained a variety of genes involved with neural processing that were differentially expressed between trained and naïve male eyes, including additional neurotransmitter receptors (acetylcholine receptor subunit alpha-like 2, gamma-aminobutyric acid type B receptor subunit 2 isoform X3, and dopamine receptor 2-like). While there is some debate over whether eyes should be considered part of the peripheral nervous system or the central nervous system in vertebrates (London et al., 2013), there has been less attention given to the potentially broad cognitive role of the retina in comparison to the optic lobe in insects (as illustrated by (Perry et al., 2017)). Our findings not only indicate that transcription in the butterfly eye changes in response to social scenario (presence/absence of a sexually mature conspecific), but that this change includes the transcription of genes associated with higher processing, indicating that neurogenomic processes associated with cognition might not be limited to the optic lobe and central brain in insects, but might also occur in the retina.

### Broad role of sensory receptors and neurotransmitters in sexually dimorphic behavior

Although neurogenomic assessment of sexually dimorphic behavior is relatively rare to date, similarities between our results and those in other animal systems suggest common mechanisms may underlie sexually dimorphic behavior across animal taxa. Sensory receptors seem to be especially important and connected to downstream sexually dimorphic gene networks. For example, odorant receptor expression influences female receptivity and male ability to differentiate between the sexes in *D. melanogaster* (Datta et al., 2008), male and female zebra finches exhibit different brain gene expression profiles when listening to the same song (Gobes et al., 2009), a number of butterfly species exhibit sexually dimorphic opsin expression patterns (Buerkle et al., 2022; Everett et al., 2012), and *B. anynana* exhibit sexually dimorphic chemical receptor expression during a mate preference learning event (this study). Sexually dimorphic catecholamine-associated expression (receptors or binding proteins, for example) also appears to be important for driving sexually dimorphic social behaviors across taxa, as illustrated by sex-dependent distribution of tyrosine hydroxylase in male and female plainfin midshipman fish brains (Goebrecht et al., 2014) and sexually dimorphic association of dopamine receptors and binding proteins with social interactions in *B. anynana* butterflies (this study). Pathways integrating sensory receptors and catecholamine neurotransmitters may be particularly fruitful for future study of sexually dimorphic behaviors across animal taxa.

### Wing patterning genes may be “magic” genes

While butterfly wing patterning genes have long been hypothesized to play a role in shaping both wing pattern and preference for wing pattern (Kronforst and Papa, 2015; Kronforst et al., 2006; Merrill et al., 2015; Merrill et al., 2019; Naisbit et al., 2001), evidence supporting this hypothesis has been rare. Here we show that a number of wing patterning genes are differentially expressed in the brain and eyes during a sexual (training) encounter. Not only are these genes associated with wing patterning in a range of butterfly species, but a subset of these genes are specifically associated with aspects of eyespot production in *B. anynana* (Brunetti et al., 2001; Ozsu and Monteiro, 2017; Prakash and Monteiro, 2018) and/or with UV reflectance (Ficarrotta et al., 2022).

Because male and female *B. anynana* learn preferences for eyespot number, and specifically the UV-reflective center of the eyespots (Westerman et al., 2012; Westerman et al., 2014), these genes that both influence eyespots or UV scale production and are differentially expressed in the brain or eyes during an intersexual social encounter (*invected*, *spalt*, *apterous*, *CD63 antigen-like*, and *bric-a-brac*) are particularly promising candidate magic genes in the *B. anynana* system. The brain and eye expression profiles of genes known to influence wing patterning traits important for mate selection in other butterfly systems, such as *BarH-1* (Woronik et al., 2019), *artistaless* (Westerman et al., 2018), *cortex* (Nadeau et al., 2016), and *doublesex* (Kunte et al., 2014), support the hypothesis that these genes may be expressed in the brains or eyes of the butterfly species using these genes to control wing pattern elements under sexual selection. Future studies should explore the pervasiveness of genes influencing both wing pattern and mate preference across the Lepidoptera.

### Conclusions

Here we show that sexually dimorphic, imprinting-like learning is associated with sexually dimorphic gene expression in the brains and eyes of *B. anynana* butterflies during a training event. Differentially expressed genes include sensory receptors and genes associated with neurotransmitters in both tissue types, indicating dynamic and sex-specific responses to social scenario in both the peripheral and central nervous systems. Sexually dimorphic expression of chemosensory genes supports the role of pheromones in female but not male imprinting-like learning, while the learning-related expression of numerous wing patterning genes highlight the potential for these genes to influence both wing pattern and mate preference. Future research should explore the gene and neural networks bridging sexually dimorphic sensory receptors to sexually dimorphic behavior, and determine the functional role of wing patterning genes in mate preference in other lepidopterans.

## Materials and Methods

### Study Species and Husbandry

*Bicyclus anynana* is a sub-tropical African butterfly that has been reared in the lab since 1988. The colony at the University of Arkansas was established in spring 2017 from ∼1,000 eggs derived from a population in Singapore. Butterflies at the University of Arkansas were reared in a climate-controlled greenhouse at ∼27°C, 70% humidity, and under a 13:11h light:dark cycle to mimic wet season conditions and ensure development of the wet season phenotype (Brakefield and Reistma, 1991). Butterflies bred in the laboratory have levels of genetic diversity comparable to those in natural populations, as suggested by similar single-nucleotide polymorphism frequencies found in laboratory and natural populations (Beldade et al., 2006; de Jong et al., 2013).

All adult butterflies used in this study hatched from eggs laid on young corn plants (*Zea mays*) in breeding colony cages containing ∼200-500 male and female *B. anynana* butterflies. Plants with eggs were moved to cages containing additional corn plants for larval consumption, and larvae were fed *ad libitum* until pupation. Upon pupation, pupae were placed in mesh cages (31.8 cm × 31.8 cm × 31.8 cm; Bioquip, Compton, CA, USA) until emergence. Upon emergence, butterflies were transferred to sex- and age-specific cages to isolate the sexes from one another. All butterflies were provided with fresh banana every other day.

### Behavioral assays and sample collection

All behavioral assays and sample collection took place between November 2018 - July 2019. Within one hour of dawn, assays were conducted by placing butterflies in a novel mesh cage (39.9 cm × 39.9 cm × 59.9 cm; Bioquip, Compton, CA, USA) for a three-hour observation period (Fig. 1A). Training behavioral assays consisted of either: (1) a newly emerged male paired with a two-day-old, zero-spot female, for which black paint (Enamel Glossy Black 1147, Testors, Rockford, IL, USA) was applied directly on top of her two dorsal hindwing eyespot pupils to block all UV reflectance (for details see (Westerman et al., 2014)) or (2) a newly emerged female paired with a two-day-old, four-spot male, for which UV-reflective paint (White, FishVision, Fargo, ND, USA) was applied between the two natural dorsal forewing eyespot pupils to create two extra eyespot pupils (for details see (Westerman et al., 2012)). The UV-reflective paint closely replicated the reflectance spectra of natural *B. anynana* eyespot pupils (Westerman et al., 2012). All eyespot manipulations were performed one day prior to behavioral watches. Control assays consisted of either one newly emerged male or one newly emerged female placed in a novel mesh cage (39.9 cm × 39.9 cm × 59.9 cm; Bioquip, Compton, CA, USA) in isolation. For any given training assay, a control assay using the same sex as the training assay focal animal was conducted concurrently (e.g., for a newly emerged male + zero-spot female training assay, a control assay consisting of a newly emerged male in isolation was run in tandem). All behaviors exhibited by the observed butterflies were recorded using SpectatorGo! (BIOBSERVE; Bonn, Germany). Observed behaviors included: *flutter, fly, walk, rest* (wings closed)*, bask* (wings open greater than 45°)*, antenna wiggle, court* (as defined in (Nieberding et al., 2008), and *copulate*.

After the three-hour behavioral watch, each butterfly’s head was removed with RNase-free scissors, transferred into a RNase-free microcentrifuge tube (Biotix; San Diego, CA, USA), and immediately flash frozen in liquid nitrogen. Frozen samples were then stored in a -80⁰C freezer until dissection and RNA extraction. We collected the heads of ten individuals per group (trained male, trained female, naïve male, and naïve female) to account for variation in response to training, as previous studies suggest that ∼75% of females and ∼80% of males learn to prefer the trainer phenotype after a three-hour training exposure (Westerman et al., 2012; Westerman et al., 2014).

### RNA extraction and cDNA library preparation

To prevent RNA degradation during processing, heads were immersed in 500 µL of pre-chilled RNAlater-ICE (Ambion; Austin, TX, USA) and incubated at -20°C for approximately 18 hours prior to dissection. Thawed heads were then dissected under a dissecting microscope (Zeiss Stemi 508; Jena, Germany) while submerged in RNAlater-ICE to isolate eye and brain tissue. The eyes and brain for each sample were mechanically disrupted separately in lysis buffer using RNase-free, disposable pestles, and small (<200 nucleotides) and large (>200 nucleotides) RNA were extracted separately for each tissue with the NucleoSpin® miRNA kit (Macherey-Nagel; Düren, Germany). RNA quality and quantity were determined using a NanoDrop 2000 (Thermo Fisher Scientific; Waltham, MA, USA), Qubit 2.0 (Invitrogen; Waltham, Massachusetts, USA), and TapeStation 2200 (Agilent; Santa Clara, CA, USA).

Libraries were prepared for the eyes (left and right eye together; n=40) and brain (n=40) for each individual using the KAPA mRNA HyperPrep Kit and Unique Dual-Indexed Adapters (KAPA Biosystems; Wilmington, MA, USA), with 100 ng of large RNA as input. After running all cDNA libraries on a TapeStation 2200 (Agilent; Santa Clara, CA, USA) and confirming that they were of high quality, libraries were shipped to the University of Chicago Genomics Facility on dry ice. All libraries were subjected to an additional quality assessment using a 5300 Fragment Analyzer (Agilent; Santa Clara, CA, USA), followed by 50 bp SE sequencing across eight lanes of a HiSeq 4000 (Illumina; San Diego, CA, USA).

### Read trimming, alignment, and quantification

We concatenated the raw fastq files from all eight lanes for each library and performed an initial quality assessment using FastQC v0.11.5 (http://www.bioinformatics.babraham.ac.uk/projects/fastqc/). One sample (TME_A3, a trained eye sample) failed to sequence properly, so was discarded from downstream analysis. Trimmomatic v0.38 was used to remove any Illumina sequencing adapters from the raw reads (Bolger et al., 2014). We then aligned the adapter-trimmed reads for each sample to the most recent *B. anynana* reference genome (v1.2; (Nowell et al., 2017)) using STAR v2.7.1a (Dobin et al., 2013) and quantified all reads using the “--quantMode GeneCounts” option, which is equivalent to counts produced by the htseq-count script from HTSeq (Anders et al., 2015).

### Differential gene expression analyses

The read counts generated by STAR were used as input for the DESeq2 v1.24.0 package (Love et al., 2014) for R (Version 3.6.2, R Foundation for Statistical Computing, Vienna, Austria) to conduct differential expression analyses. Specifically, we used the generalized linear model design:

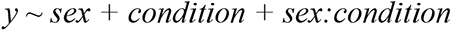

where expression (*y*) is a function of *sex* (male or female), *condition* (trained or naïve), and their interaction (*sex:condition*). With this design, we made five different tissue-specific comparisons: (1) naïve females vs. naïve males; (2) trained females vs. trained males; (3) trained females vs. naïve females; (4) trained males vs. naïve males; and (5) the interaction of sex and condition. To investigate the overall effect of training while controlling for differences in expression specific to sex, we performed an additional tissue-specific analysis that utilized the design:

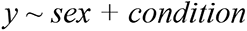

Only genes with ≥10 total mapped reads were used for the differential expression analyses. Gene expression comparisons were conveyed as the binary log of the expression fold change (log_2_FC), with log_2_FC shrinkage performed using the ashr method (Stephens, 2017) to obtain more accurate estimates of effect size. Genes were considered differentially expressed if they had a false discovery rate (FDR; (Benjamini and Hochberg, 1995)) < 0.05.

In addition to these standard differential expression analyses, we also performed permutation tests similar to those utilized in Ghalambor et al. (2015) and Bloch et al. (2018). Because this method does not assume gene independence (an unlikely assumption given the nature and abundance of gene co-expression networks), the risk of over-correction is reduced compared to other multiple testing correction methods, resulting in a more accurate representation of the expression data structure (Slonim, 2002). For each tissue, we randomly assigned both the sex and treatment for each sample to create 1,000 permuted sample phenotype tables. For each of the reassigned sample sets, we ran the DESeq2 analysis exactly as we had for the original analysis, ultimately resulting in a null distribution of 1,000 p-values for every gene. For any given gene, if the p-value from the original analysis was less than the 1% tail of the permuted null distribution, it was considered differentially expressed. Annotations for all differentially expressed genes, including the identified putative vision-and chemsensory-related gene annotations, were extracted from the *B. anynana* reference genome functional annotation from (Ernst and Westerman, 2021).

### Weighted gene co-expression network analyses

We performed separate weighted gene co-expression network analyses (WGCNA) for the brain and eyes using the WGCNA v1.70-3 R package (Langfelder and Horvath, 2008) following the WGCNA package developers’ recommendations. We first preprocessed the expression data by removing all genes with <10 reads in >90% of the samples to minimize noise from lowly-expressed genes, and a variance-stabilizing transformation was performed on the remaining data using the “varianceStabilizingTransformation” function in DESeq2. Signed co-expression networks for each tissue were constructed by building an adjacency matrix with type = “signed,” topological overlap matrix (TOM) with TOMType = “signed,” and the soft-thresholding power set to 12 for brains and 14 for eyes. We then identified modules of co-expressed genes using the “cutreeDynamic” function with the following parameters: deepSplit = 2, pamRespectsDendro = FALSE, and minClusterSize = 30. After initial module identification, we merged modules of high co-expression similarity by first calculating and clustering their eigengenes (the first principal component of a module representing its gene expression profile (Langfelder and Horvath, 2008)) and employing the “mergeCloseModules” function with the “cutHeight” set to 0.25.

To identify modules that were significantly associated with any of the sample traits, we used the “binarizeCategoricalVariable” function to create pairwise binary indicators (“traits”) for our contrasts of interest (i.e., naïve male vs. naïve female, trained female vs. naïve female, trained male vs. naïve male, and trained female vs. naïve female) and correlated eigengenes with these sample traits. We then adjusted all p-values using the FDR method (Benjamini and Hochberg, 1995), and any module-trait correlations with an FDR <0.05 were considered significant. For all modules that showed significant associations with sample traits, hub genes (genes with the highest connectivity) were identified using the “chooseTopHubInEachModule” function.

For visualization and further analysis, both networks were then exported to Cytoscape v3.8.2 (Shannon et al., 2003) using the “exportNetworkToCytoscape” function with “threshold” set to 0.02. The Cytoscape “Network Analyzer” tool was used to obtain further statistics regarding the connectivity of genes within the network. Specifically, we calculated three statistics for each gene: (1) degree (the number of other genes connected to a given gene, with a larger number indicating a more highly connected gene), (2) neighborhood connectivity (the average connectivity of all of a gene’s neighboring genes), and (3) clustering coefficient (how connected a gene is to its neighboring genes relative to how connected it could be, with “0” representing completely unconnected and “1” representing maximum connectivity).

### Gene Ontology Analyses

To facilitate the characterization of DEG sets and significant modules, GO enrichment analyses were performed using the Fisher’s Exact Test function in Blast2GO v5.2.5 (Conesa et al., 2005) with the GO annotations extracted from Ernst and Westerman (2021). In each case, all genes in the expression set (for the WGCNA analysis, all genes that were used in the co-expression analysis) for the respective tissue were used as the reference set, and an FDR threshold of <0.05 was set to identify significantly enriched GO terms. All DEG sets and significant modules were tested for GO enrichment.

To further explore the differences between male and female tissues for each condition, we used GOExpress v1.20.0 (Rue-Albrecht et al., 2016) to identify GO terms that best classify the samples from two groups (e.g., female trained brains and male trained brains) based on their gene expression profiles. For these analyses, reads were first normalized to counts per million (CPM) with edgeR v3.28.1 (Robinson et al., 2010), and only genes with ≥1 CPM for at least 10 samples (the maximum number of replicates per group) were retained for the input expression matrix. The random forest was set to 10,000 trees, and GO terms that were associated with at least five genes and with a p-value <0.05 after 1,000 permutations were considered significant.

### Identification of wing patterning genes

In addition to examining differential expression, co-expression networks, and GO signatures, we also investigated the expression patterns of known wing patterning genes, as these genes have been hypothesized to act as “magic genes” and to have the capacity to influence both preference as well as the preferred trait (Servedio, 2009; Smadja and Butlin, 2011; Westerman, 2019). Specifically, we used the functional annotations and butterfly wing patterning gene list from Ernst and Westerman (2021) to identify wing patterning genes expressed in eye and brain tissue and to determine if they were differentially expressed between the sexes. The genes included numerous *B. anynana* wing patterning genes (Beldade and Peralta, 2017; Bhardwaj et al., 2018; Connahs et al., 2019; Matsuoka and Monteiro, 2018; Monteiro et al., 2013; Monteiro et al., 2006; Monteiro and Prudic, 2010; Ozsu et al., 2017; Prakash and Monteiro, 2018, 2020; Saenko et al., 2011), as well as genes characterized in other butterfly species (Ficarrotta et al., 2022; Martin and Reed, 2010; Nadeau et al., 2016; Reed et al., 2011; Westerman et al., 2018; Woronik et al., 2019).

### Analysis of Behavior

We first conducted a Shapiro-Wilk test to assess normality of the behavioral data. We then performed a Kruskal-Wallis test to examine the effect of sex on behavior, followed by a second Kruskal-Wallis test subset by treatment (naïve, trained, trainer) to test for the effect of sex on behavior in each treatment. We conducted a principal components analysis (PCA) on behavior to search for hidden correlations and create new composite variables (S39 Table). We then performed a Kruskal-Wallis test to test for the effect of sex on PC1, PC2, and PC3. We calculated a Bonferroni correction to account for multiple testing, producing an adjusted significance value of p = 0.0025.

### Ethical Note

All *B. anynana* butterflies were maintained in laboratory conditions as specified by U.S. Department of Agriculture APHIS permit P526P-17-00343. Butterflies not used for this experiment were maintained with ample food and water until natural death.

## Acknowledgements

We thank Matthew Murphy, Grace Hirzel, Deonna Robertson, and Sushant Potdar for assistance with animal husbandry. This research was supported by NSF IOS grant #1937201 to ELW, an Arkansas Biosciences Institute (the major research component of the Arkansas Tobacco Settlement Proceeds Act of 2000) grant to ELW, a University of Arkansas Honors College grant to GAA & ELW, the Arkansas High Performance Computing Center, which is funded through multiple NSF grants and the Arkansas Economic Development Commission, and the University of Arkansas.

## Data Availability

All raw sequence data associated with this study are accessible through the NCBI Sequence Read Archive (SRA) database under XXX. Behavioral data are available at Dryad Database XXXX. All other data presented in this study are available within this manuscript and its Supplemental Information.

