## Supplemental Information for "A learning experience elicits sex-dependent neurogenomic responses in *Bicyclus anynana* butterflies"

**S1 Figure:** **PCA plot of PC1 and PC2 based on variance stabilized counts for all samples**. Both brain samples and eye samples cluster separately from one another, with the exception of one sample (TMB_E2). Given that tissue type explains approximately 85% of the variance, we conclude that this sample was likely mislabeled and was thus excluded from downstream analyses.

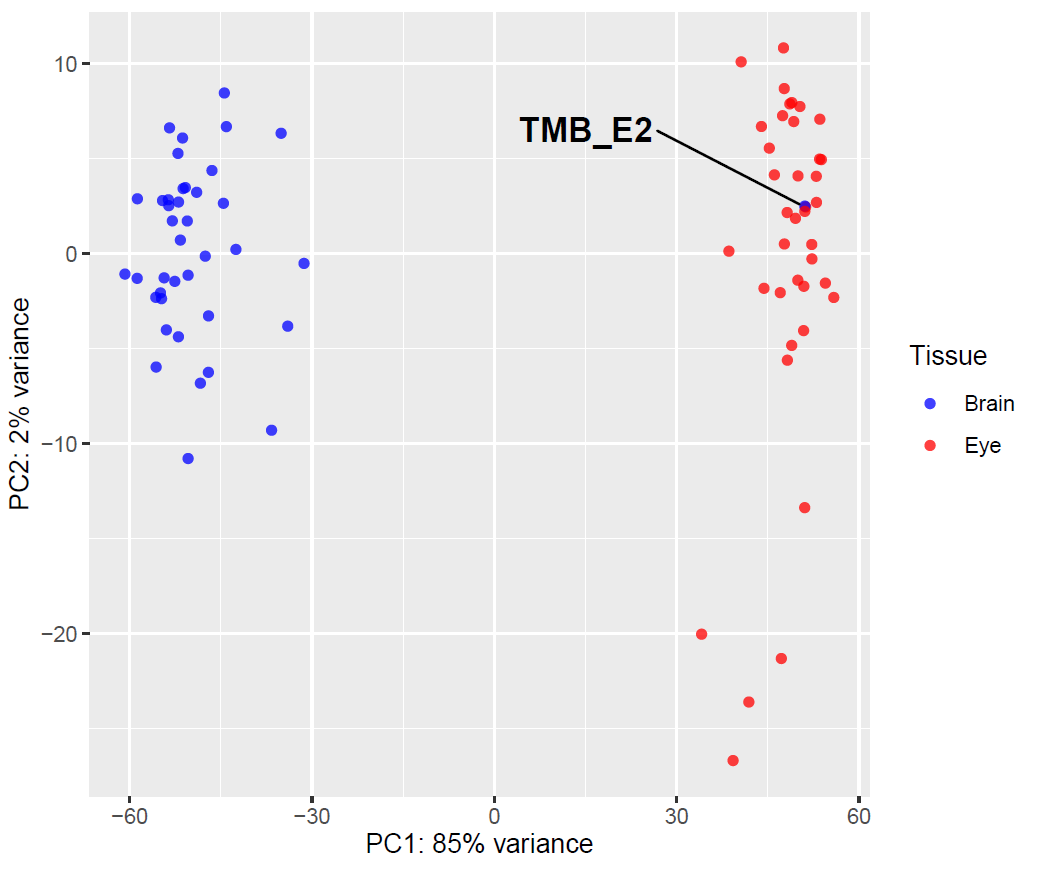

**S2 Figure:** **Gene expression heatmaps of differentially expressed genes from the sex-independent trained vs. naïve comparisons.** A) Trained brains vs. naïve brains comparison. Each row indicates a single gene, and each column indicates an individual sample. Counts were first normalized by variance stabilizing transformation, and gene-wise Z-scores were calculated for plotting. Genes and samples are clustered by expression, with warmer colors denoting increased expression relative to the mean for a given gene, while cooler colors denote decreased expression relative to the mean. B) Trained eyes vs. naïve eyes comparison.

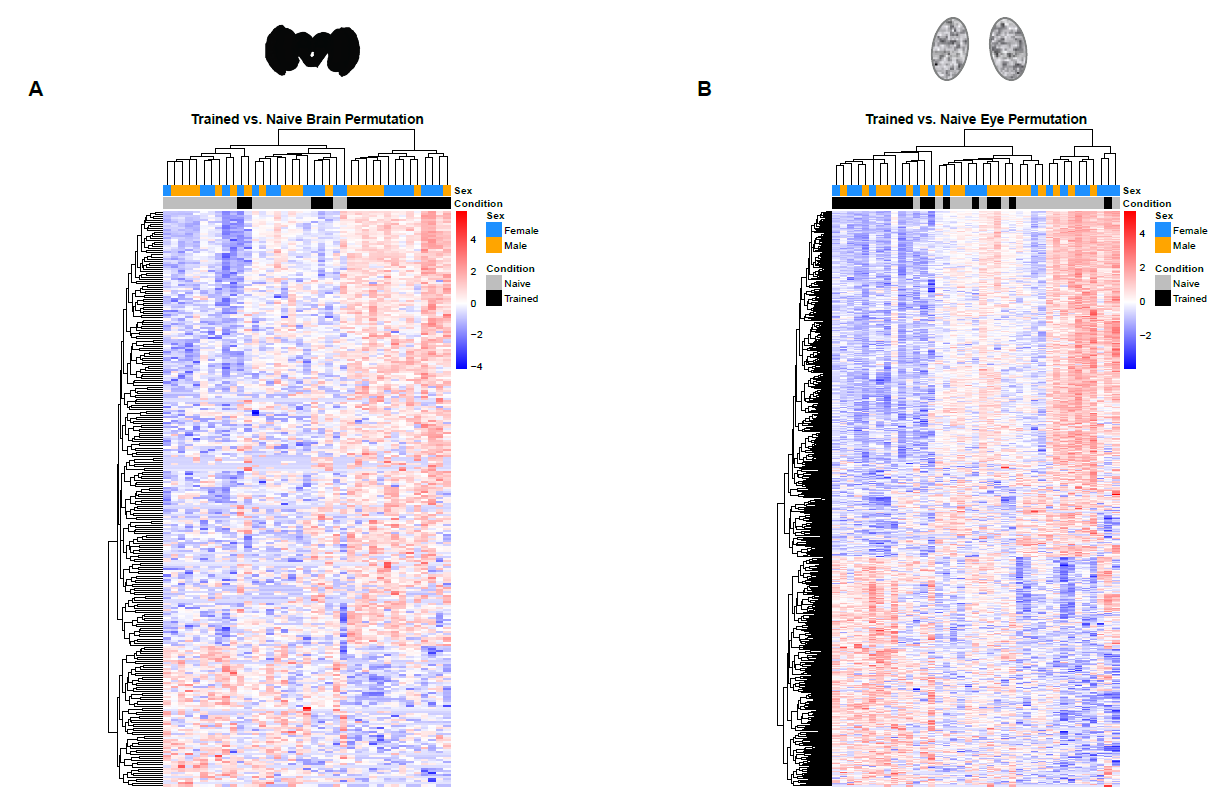

**S3 Figure:** **WGCNA module clustering and significant modules for brains**. A) Gene clustering dendrogram for brains. All identified co-expression modules are represented by the colors in the “Dynamic Tree Cut” bar, while the colors in the “Merged dynamic” bar represent the final modules after merging highly similar modules. B) Cytoscape plot of the red module for the WGCNA brain analysis. Each circle indicates a node (gene), and each grey line indicates an edge (connection). The size of each circle represents its degree (connectivity) within the module, with larger circles denoting more highly connected genes. Circles highlighted with color indicate genes that were differentially expressed from the permutation analysis in the comparison that was significantly associated with the module, whereas open circles denote the rest of the genes in that module. TMB_NMB_perm = differentially expressed in the trained male brain vs. naïve male brain permutation analysis; Not_DE = not differentially expressed (i.e., all other genes).

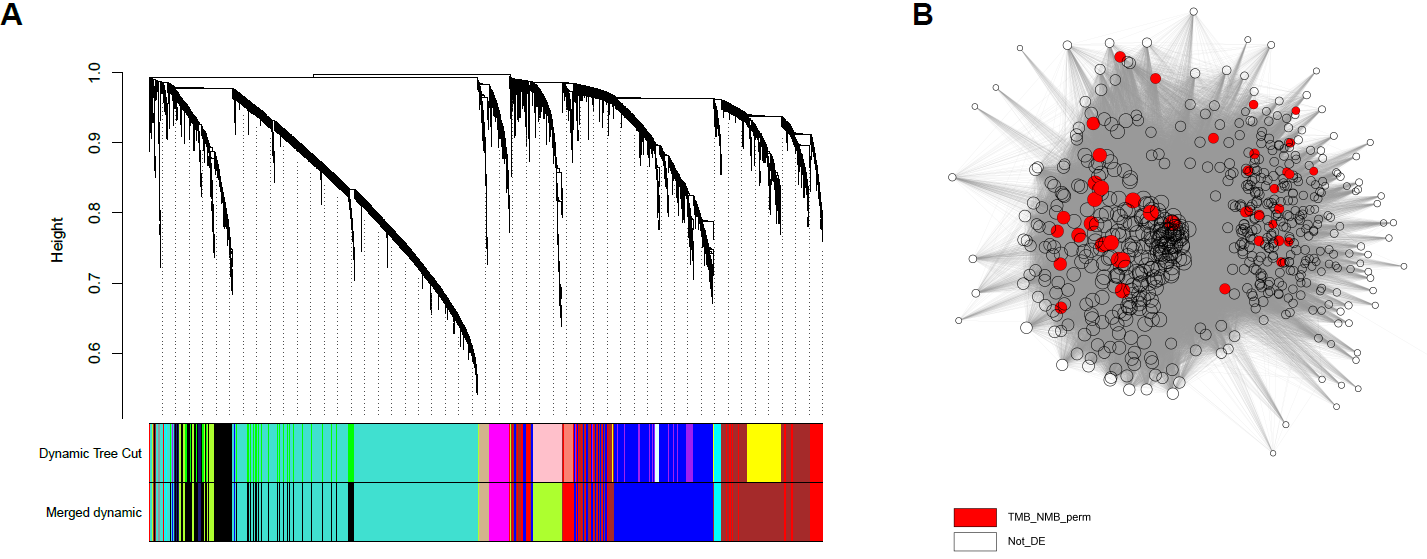

**S4 Figure:** **WGCNA module clustering and significant modules for eyes.** A) WGCNA gene clustering dendrogram for eyes. All identified co-expression modules are represented by the colors in the “Dynamic Tree Cut” bar, while the colors in the “Merged dynamic” bar represent the final modules after merging highly similar modules. B) Cytoscape plot of the grey60 module for the WGCNA eye analysis. Each circle indicates a node (gene), and each grey line indicates an edge (connection). The size of each circle represents its degree (connectivity) within the module, with larger circles denoting more highly connected genes. Circles highlighted with color indicate genes that were differentially expressed from the permutation analysis in the comparisons that were significantly associated with the module, whereas open circles denote the rest of the genes in that module. C) Cytoscape plot of the black module for the WGCNA eye analysis. D) Cytoscape plot of the magenta module for the WGCNA eye analysis. E) Cytoscape plot of the blue module for the WGCNA eye analysis. F) Cytoscape plot of the cyan module for the WGCNA eye analysis. G) Cytoscape plot of the tan module for the WGCNA eye analysis. H) Cytoscape plot of the midnightblue module for the WGCNA eye analysis. TME_NME_perm = differentially expressed in the trained male eye vs. naïve male eye permutation analysis; TME_NME_perm & NFE_NME_perm = differentially expressed in both the trained male eye vs. naïve name eye and naïve female eye vs. naïve male eye permutation analyses; NFE_NME_perm = differentially expressed in the naïve female eye vs. naïve male eye permutation analysis; TFE_NFE_perm = differentially expressed in the trained female eye vs. naïve female eye permutation analysis; Not_DE = not differentially expressed (i.e., all other genes).

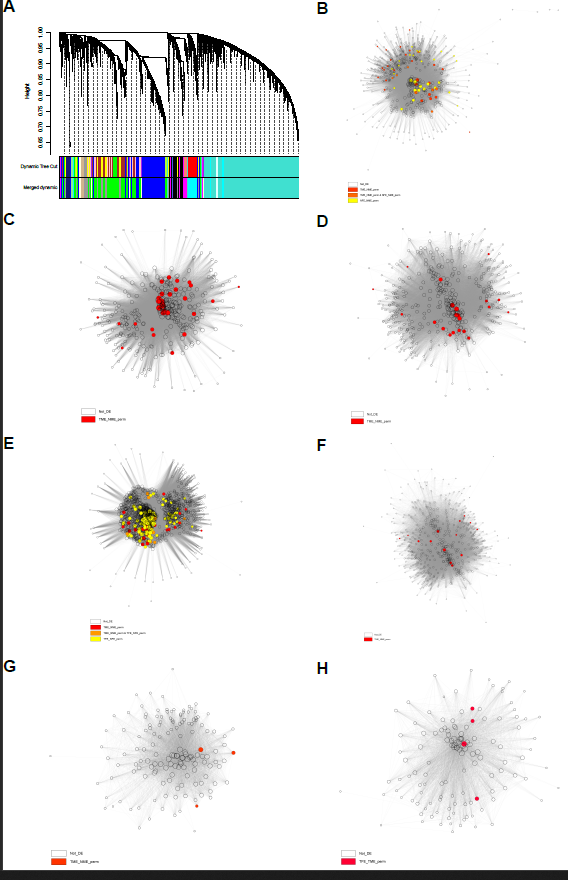

**Legends for Supplemental Tables, found in associated excel file.**

**S1 Table: No effect of sex on composite behaviors (PC1, PC2, PC3)** during isolation (naïve), when interacting with a young butterfly of the opposite sex( trainer), or being exposed to a sexually mature butterfly of the opposite sex (trained).

**S2 Table: No effect of sex on behavior** during isolation (naïve), when interacting with a young butterfly of the opposite sex( trainer), or being exposed to a sexually mature butterfly of the opposite sex (trained).

**S3 Table:** Summary statistics for sequenced libraries. The Mean Mapped Reads column contains the mean number of reads per sample that mapped to the B. anynana reference genome (v1.2). The Mean Reads for DE column contains the mean number of reads per sample that aligned to genes that were overlapped by ≥10 reads across all tissue-specific libraries (e.g., across all brain libraries for the brain analysis) and were subsequently used for differential expression analysis. All reads statistics are shown in millions and exclude sample TMB_E2, which was determined to be mislabeled based on clustering analysis.

**S4 Table: Genes differentially expressed between naïve female and naïve male brains.** Perm_Threshold indicates the 1% p-value permutation threshold and DE_in_Standard_DESeq2 and DE_in_Perm_DESeq2 indicate whether the gene was differentially expressed in the standard or permutation-based DESeq2 analysis, respectively.

**S5 Table: Genes differentially expressed between trained female and trained male brains.** Perm_Threshold indicates the 1% p-value permutation threshold and DE_in_Standard_DESeq2 and DE_in_Perm_DESeq2 indicate whether the gene was differentially expressed in the standard or permutation-based DESeq2 analysis, respectively. Unique_to_Training_Perm indicates whether differentially expressed genes from the permutation analysis are also differentially expressed in the naive comparison.

**S6 Table: Genes differentially expressed between naïve female and naïve male eyes.** Perm_Threshold indicates the 1% p-value permutation threshold and DE_in_Standard_DESeq2 and DE_in_Perm_DESeq2 indicate whether the gene was differentially expressed in the standard or permutation-based DESeq2 analysis, respectively.

**S7 Table: Genes differentially expressed between trained female and trained male eyes.** Perm_Threshold indicates the 1% p-value permutation threshold and DE_in_Standard_DESeq2 and DE_in_Perm_DESeq2 indicate whether the gene was differentially expressed in the standard or permutation-based DESeq2 analysis, respectively. Unique_to_Training_Perm indicates whether differentially expressed genes from the permutation analysis are also differentially expressed in the naive comparison.

**S8 Table: Genes differentially expressed between trained and naïve female brains.** Perm_Threshold indicates the 1% p-value permutation threshold and DE_in_Standard_DESeq2 and DE_in_Perm_DESeq2 indicate whether the gene was differentially expressed in the standard or permutation-based DESeq2 analysis, respectively.

**S9 Table: Genes differentially expressed between trained and naïve female eyes.** Perm_Threshold indicates the 1% p-value permutation threshold and DE_in_Standard_DESeq2 and DE_in_Perm_DESeq2 indicate whether the gene was differentially expressed in the standard or permutation-based DESeq2 analysis, respectively.

**S10 Table: Genes differentially expressed between trained and naïve male brains.** Perm_Threshold indicates the 1% p-value permutation threshold and DE_in_Standard_DESeq2 and DE_in_Perm_DESeq2 indicate whether the gene was differentially expressed in the standard or permutation-based DESeq2 analysis, respectively.

**S11 Table: Genes differentially expressed between trained and naïve male eyes**. Perm_Threshold indicates the 1% p-value permutation threshold and DE_in_Standard_DESeq2 and DE_in_Perm_DESeq2 indicate whether the gene was differentially expressed in the standard or permutation-based DESeq2 analysis, respectively.

**S12 Table: Genes with a significant sex:condition interaction for brains.** Perm_Threshold indicates the 1% p-value permutation threshold and DE_in_Standard_DESeq2 and DE_in_Perm_DESeq2 indicate whether the gene was differentially expressed in the standard or permutation-based DESeq2 analysis, respectively.

**S13 Table: Genes with a significant sex:condition interaction for eyes**. Perm_Threshold indicates the 1% p-value permutation threshold and DE_in_Standard_DESeq2 and DE_in_Perm_DESeq2 indicate whether the gene was differentially expressed in the standard or permutation-based DESeq2 analysis, respectively.

**S14 Table: Genes differentially expressed between trained and naïve brains, independent of sex.** Perm_Threshold indicates the 1% p-value permutation threshold and DE_in_Standard_DESeq2 and DE_in_Perm_DESeq2 indicate whether the gene was differentially expressed in the standard or permutation-based DESeq2 analysis, respectively.

**S15 Table: Genes differentially expressed between trained and naïve eyes, independent of sex**. Perm_Threshold indicates the 1% p-value permutation threshold and DE_in_Standard_DESeq2 and DE_in_Perm_DESeq2 indicate whether the gene was differentially expressed in the standard or permutation-based DESeq2 analysis, respectively.

**S16 Table**: Genes of interest for the various differential expression analyses.

**S17 Table: GO terms significantly associated with differences between naïve female and naïve male brains from GOExpress analysis**. Average_Rank and Average_Score denote the respective means for all genes that were annotated with a given GO term. Total_Count indicates the number of genes in the annotation assigned to each GO term, and Data_Count indicates the number of genes in the GOExpress ExpressionSet assigned to each GO term.

**S18 Table: GO terms significantly associated with differences between trained female and trained male brains from GOExpress analysis**. Average_Rank and Average_Score denote the respective means for all genes that were annotated with a given GO term. Total_Count indicates the number of genes in the annotation assigned to each GO term, and Data_Count indicates the number of genes in the GOExpress ExpressionSet assigned to each GO term. Unique_to_Training indicates significant GO terms specific to differences between trained female and trained male brains.

**S19 Table: GO terms significantly associated with differences between naïve female and naïve male eyes from GOExpress analysis**. Average_Rank and Average_Score denote the respective means for all genes that were annotated with a given GO term. Total_Count indicates the number of genes in the annotation assigned to each GO term, and Data_Count indicates the number of genes in the GOExpress ExpressionSet assigned to each GO term.

**S20 Table: GO terms significantly associated with differences between trained female and trained male eyes from GOExpress analysis**. Average_Rank and Average_Score denote the respective means for all genes that were annotated with a given GO term. Total_Count indicates the number of genes in the annotation assigned to each GO term, and Data_Count indicates the number of genes in the GOExpress ExpressionSet assigned to each GO term. Unique_to_Training indicates significant GO terms specific to differences between trained female and trained male eyes.

**S21 Table: Enriched GO terms from Blast2GO for the trained vs. naïve female eye contrast.**

**S22 Table: Enriched GO terms for genes differentially expressed between trained and naïve eyes, independent of sex**, for the permutation analysis.

**S23 Table: Cytoscape network statistics for the red module from the brain WGCNA analysis.** Only genes that exceeded the adjacency threshold of 0.02 were included in the analysis.

**S24 Table: Enriched GO terms for genes in the red module for the brain WGCNA analysis.**

**S25 Table: Cytoscape network statistics for the grey60 module from the eye WGCNA analysis.** Only genes that exceeded the adjacency threshold of 0.02 were included in the analysis.

**S26 Table: Enriched GO terms for genes in the grey60 module for the eye WGCNA analysis.**

**S27 Table: Cytoscape network statistics for the black module from the eye WGCNA analysis.** Only genes that exceeded the adjacency threshold of 0.02 were included in the analysis.

**S28 Table: Enriched GO terms for genes in the black module for the eye WGCNA analysis.**

**S29 Table: Cytoscape network statistics for the magenta module from the eye WGCNA analysis.** Only genes that exceeded the adjacency threshold of 0.02 were included in the analysis.

**S30 Table: Enriched GO terms for genes in the magenta module for the eye WGCNA analysis.**

**S31 Table: Cytoscape network statistics for the blue module from the eye WGCNA analysis.** Only genes that exceeded the adjacency threshold of 0.02 were included in the analysis.

**S32 Table: Enriched GO terms for genes in the blue module for the eye WGCNA analysis.**

**S33 Table: Cytoscape network statistics for the cyan module from the eye WGCNA analysis.** Only genes that exceeded the adjacency threshold of 0.02 were included in the analysis.

**S34 Table: Enriched GO terms for genes in the cyan module for the eye WGCNA analysis.**

**S35 Table: Cytoscape network statistics for the tan module from the eye WGCNA analysis.** Only genes that exceeded the adjacency threshold of 0.02 were included in the analysis.

**S36 Table: Cytoscape network statistics for the midnightblue module from the eye WGCNA analysis.** Only genes that exceeded the adjacency threshold of 0.02 were included in the analysis.

**S37 Table: Enriched GO terms for genes in the midnightblue module for the eye WGCNA analysis.**

**S38 Table: Wing patterning genes expressed in *B. anynana* brains and eyes**.

**S39 Table: Principle Components Analysis Loadings for PCA** containing all behaviors exhibited by males and females during the 3hr training/isolation period immediately prior to decapitation for brain and eye RNA extraction.
